# Detecting the ecological footprint of selection

**DOI:** 10.1101/2021.05.11.442553

**Authors:** Juliette Luiselli, Isaac Overcast, Andrew Rominger, Megan Ruffley, Hélène Morlon, James Rosindell

## Abstract

The structure of communities is influenced by many ecological and evolutionary processes, but the way this manifests in classic biodiversity patterns often remains unclear. Here, we aim to distinguish the ecological footprint of selection – through competition or environmental filtering – from that of neutral processes that are invariant to species identity. We build on existing Massive Eco-evolutionary Synthesis Simulations (MESS), which uses information from three biodiversity axes – species abundances, genetic diversity, and trait variation – to distinguish between mechanistic processes. In order to correctly detect and characterise competition, we add a new and more realistic form of competition that explicitly compares the traits of each pair of individuals. Our results are qualitatively different to those of previous work in which competition is based on the distance of each individual’s trait to the community mean. We find that our new form of competition is easier to identify in empirical data compared to the alternatives. This is especially true when trait data are available and used in the inference procedure. Our findings hint that signatures in empirical data previously attributed to neutrality may in fact be the result of pairwise-acting selective forces. We conclude that gathering more different types of data, together with more advanced mechanistic models and inference as done here, could be the key to unravelling the mechanisms of community assembly.

## 1 Introduction

Understanding the assembly of ecological communities is a key goal of research in both ecology and evolution. Some studies characterise community assembly as either neutral, where individual species identities are interchangeable [1], or under selection (*sensu* Vellend [2]), where species identities have influence, for example through abiotic conditions or biotic interactions [3–6]. Such selective interactions may have varying strengths, building a continuum from neutrality (no selection) to strong selection [7]. The type and strength of species’ interactions has been shown to influence the evolution of species richness [8,9], and the ecological ramifications of evolving traits [10]. Despite recent advances, it remains challenging to characterise selection from empirical data, leading to varied opinions and conclusions. The complexity of natural ecological communities is such that unravelling the role of selection from empirical data is a formidable and unsolved computational challenge.

The question of whether competition among species is important for structuring ecological communities has been a matter of particular ongoing debate [4, 6, 11, 12]. Many studies support the idea that competition for limiting resources is the driving factor of niche differentiation, which facilitates coexistence of different species due to a high intra-specific competition, also known as density-dependence [3, 4, 13]. These niche-based competitive interactions are thought to be mediated by organismal traits [4, 14]. Yet, detecting such competition statistically, and therefore understanding its generality across systems, remains a challenge [4, 15, 16]. In contrast, neutral theory, as the prevailing alternative model to niche-based competition, is much easier to test statistically because it is a low-complexity model [17], but it is unclear whether tests that reject or fail to reject neutrality do so for valid reasons [18–20], or whether false positives or false negatives prevail.

Being able to retrieve the strength and nature of ecological competition from empirical data would be valuable to improve our understanding of competitive interactions, in ecology (shorter timescales and individual interactions) as well as in evolution (longer timescale and species interactions). One of the reasons why this has proved elusive may be that only limited data of a few types have been used to compare model predictions to reality. Multiple complementary data axes should provide more inference potential [18]. To date, competition and neutrality have largely been evaluated using species abundance distributions (SAD), as this data is historically the easiest to collect [1, 3, 19]. Other data have been used including phylogenies, which account for the evolutionary history of the local species and their past interactions [21–23], metabarcoding data, which gather abundances and genomic proximity information [24], a combination of genetic data and SADs [25, 26], and traits, which can inform on the interactions between the species and with their environment [4, 14, 27, 28]. Yet, these data are generally used in isolation from each other.

The Massive Eco-evolutionary Synthesis Simulations (MESS) model of Overcast et al. (2021) [29] allows testing mechanistic hypotheses across a combination of three data axes: species abundances, population genetic variation and trait values. These three axes reflect a variety of processes operating over a variety of time scales, from a few generations (abundances) to several tens of thousands of generations (genetic variation). Moreover, traits and genetic variation can reflect the information present in phylogenetic data, whilst the SAD and some genetic variability can be recovered from metabarcoding data. The three highlighted data axes cover the readily available and collectable data for many systems. MESS is a simulation model that can be fitted to empirical data using machine learning procedures, and thus is an ideal tool to study the eventual traces of selection in community assembly data.

Selection in the MESS model, consistent with conventional thinking [4, 14], is driven by evolving traits and interactions of individuals either with the environment or with other individuals. However, an individual’s fitness in the competition model of MESS is determined by the distance of its trait to the mean trait value of individuals in the local community, a decision made for computational convenience rather than to reflect any real mechanistic connection to the community mean trait. This “mean competition” is attractive because it delivers substantial computational gains, which are important to run enough simulations for machine learning based inference from data. Mean competition is often used to model the probability of persistence of a species [27] and has the advantage of modelling a diffuse biotic background, with which individuals interact [30]. It is, however, a weak approximation for the mechanistic reality where competition is fundamentally driven by interactions between individual organisms [31, 32]. Simulating mean competition may thus generate patterns that do not reflect real competitive processes, and so the generated data would not be appropriate for detecting competition in empirical data.

In this manuscript, we investigate the importance of competition in community assembly and our ability to detect it from empirical data through simulation models. To do this, we apply a new and more realistic pairwise competition model to the MESS system, enabled by substantial computational optimisations in the simulation method. We find that previous conclusions about the presence and strength of selection may be artefacts of the mean competition simulation method. We also find, consistent with intuition, that more data types enhance the power of inference. We show that trait data are most helpful in detection of selective forces as an alternative to neutral ones and are therefore crucial to study ecological and evolutionary forces.

## 2 Results

Community assembly model simulations progressively differentiate themselves into clusters on a PCA of summary statistics so the underlying community assembly model is easier to discriminate in older communities (Fig. 1). Results from the *β-*competition data are broadly spread across the first two PCA axes, and especially hard to distinguish from pairwise competition. However, the first two PCA components only account for around 30% of the variance, hinting that there is much more variability to be recovered elsewhere. The groups formed by pairwise competition and *β*-competition partially overlap with the neutral simulation group (Fig. 1). The filtering and mean competition groups resemble one another before reaching equilibrium (Λ < 1).

**Fig 1.**
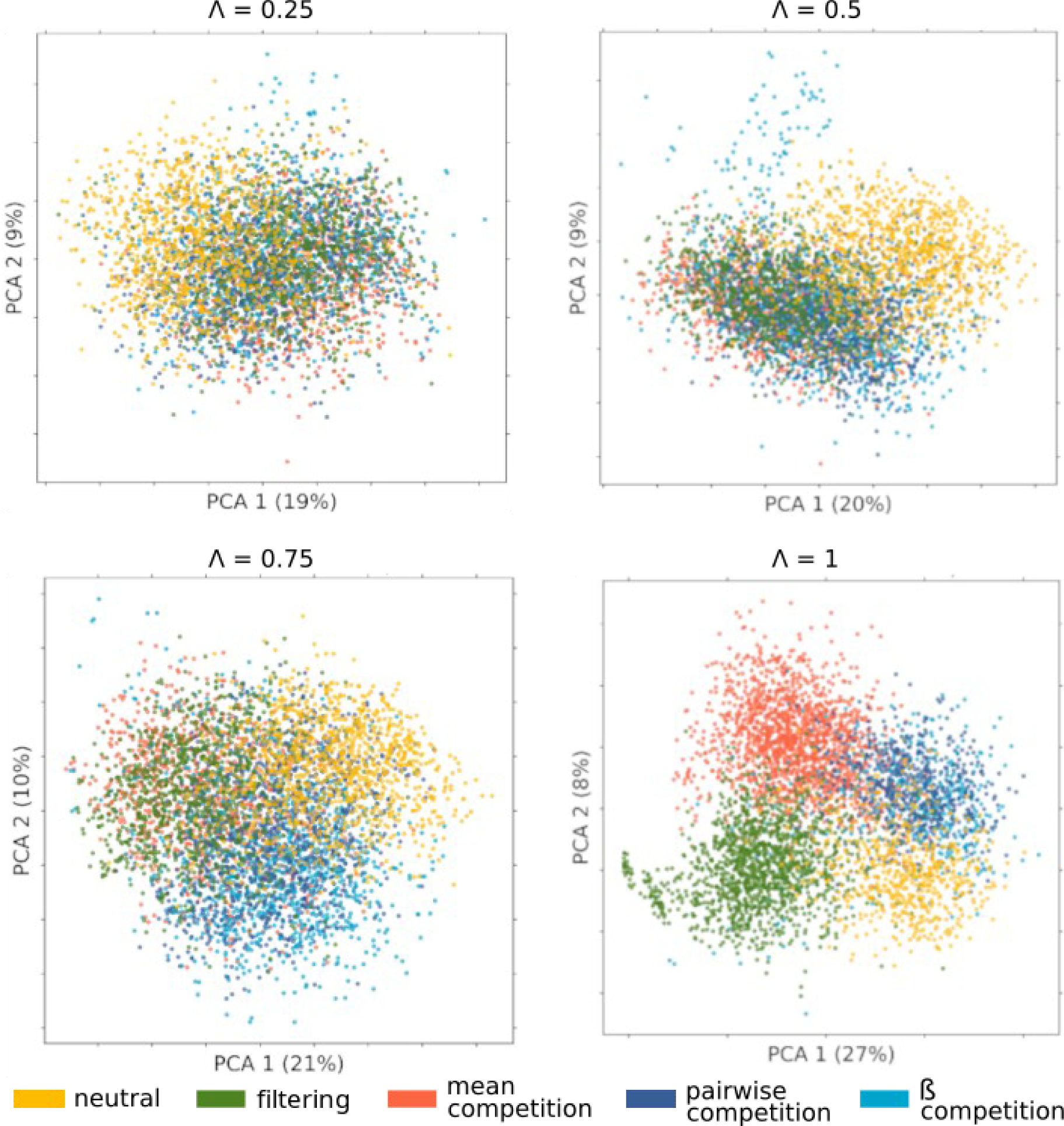
The first two principal components of the simulation summary statistics at different equilibrium stages (Λ). The different community assembly models shown as neutral (dark blue), environmental filtering (green), mean competition (orange), pairwise competition (yellow) and *β*-competition (light blue). The percentage of variance explained is indicated for each component.

Consistent results are found in the temporal dynamics of the individual summary statistics over time (Fig. 2): the summary statistics from the mean competition and environmental filtering simulations most often follow similar trajectories. The *β*-competition and pairwise competition simulations were also similar to each other (but distinct from mean competition and environmental filtering). The neutral simulations most closely resembled the *β*-competition and pairwise competition simulations (Fig. 2).

**Fig 2.**
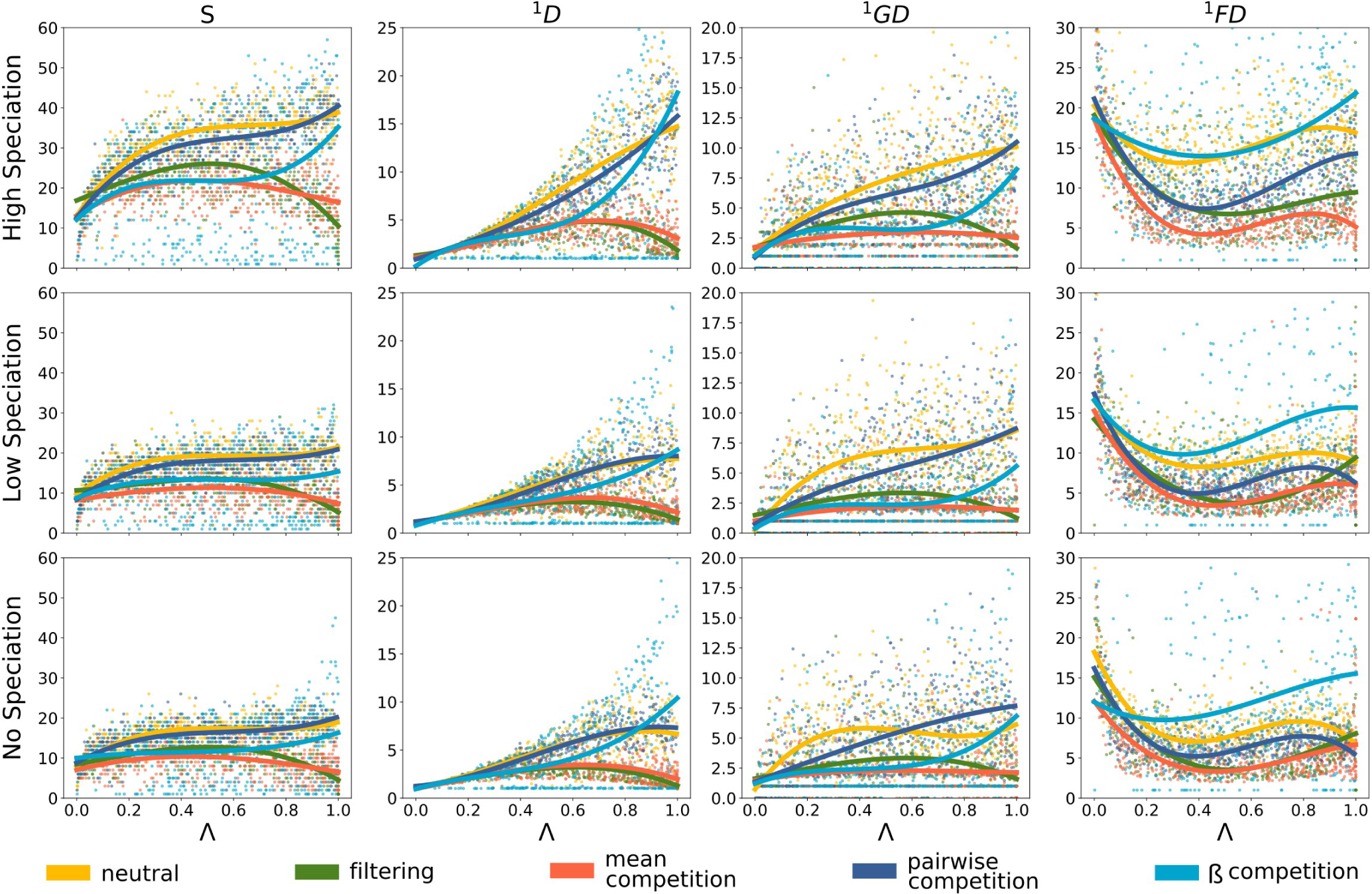
Selected community summary statistics through time for the five different community assembly models. Each panel shows a summary statistic computed at equally spaced time points for over 1500 simulations for each model, with a community size J = 1000, an ecological strength sE = 0.1 and a migration rate. Each row of panels corresponds to a different simulated speciation rate: No (ν = 0), Low (ν = 0.0005) and High (ν = 0.005). The different community assembly models are shown in the same colours as Fig. 1. Simulated values are depicted as points with a least square polynomial fitted for each community assemble model using the poly fit function of NumPy v.19.0 [33] to illustrate trajectory. The far left column of panels illustrate species richness on the y-axis (S). The y-axes of the other columns illustrate the Hill number of order 1 for abundance, genetic diversity, and trait values, respectively.

The misclassification rates when using trait values and genetic diversity show that community assembly model can be correctly determined from the simulation results in around 50% of the cases, while a random classifier would only be correct in 20% of the cases (see Fig. 3). The greatest confusion in the classifier is between pairwise competition and *β*-competition, which is expected as *β*-competition is a generalisation of pairwise competition with additional parameters. The neutral model was the best recovered by the classifier, but filtering and mean competition models were also easily distinguished by the inference procedure. A confusion matrix with SAD and genetic diversity data shows similar results (See S4 Fig. 6). The best classification is achieved when all three data types are used (See S4 Fig. 7), but the combination of all three are not yet available for empirical communities. Given the difficulty of the classifier to distinguish between pairwise competition and *β*-competition, we consider both together as an indistinguishable whole for the remainder of our analyses.

**Fig 3.**
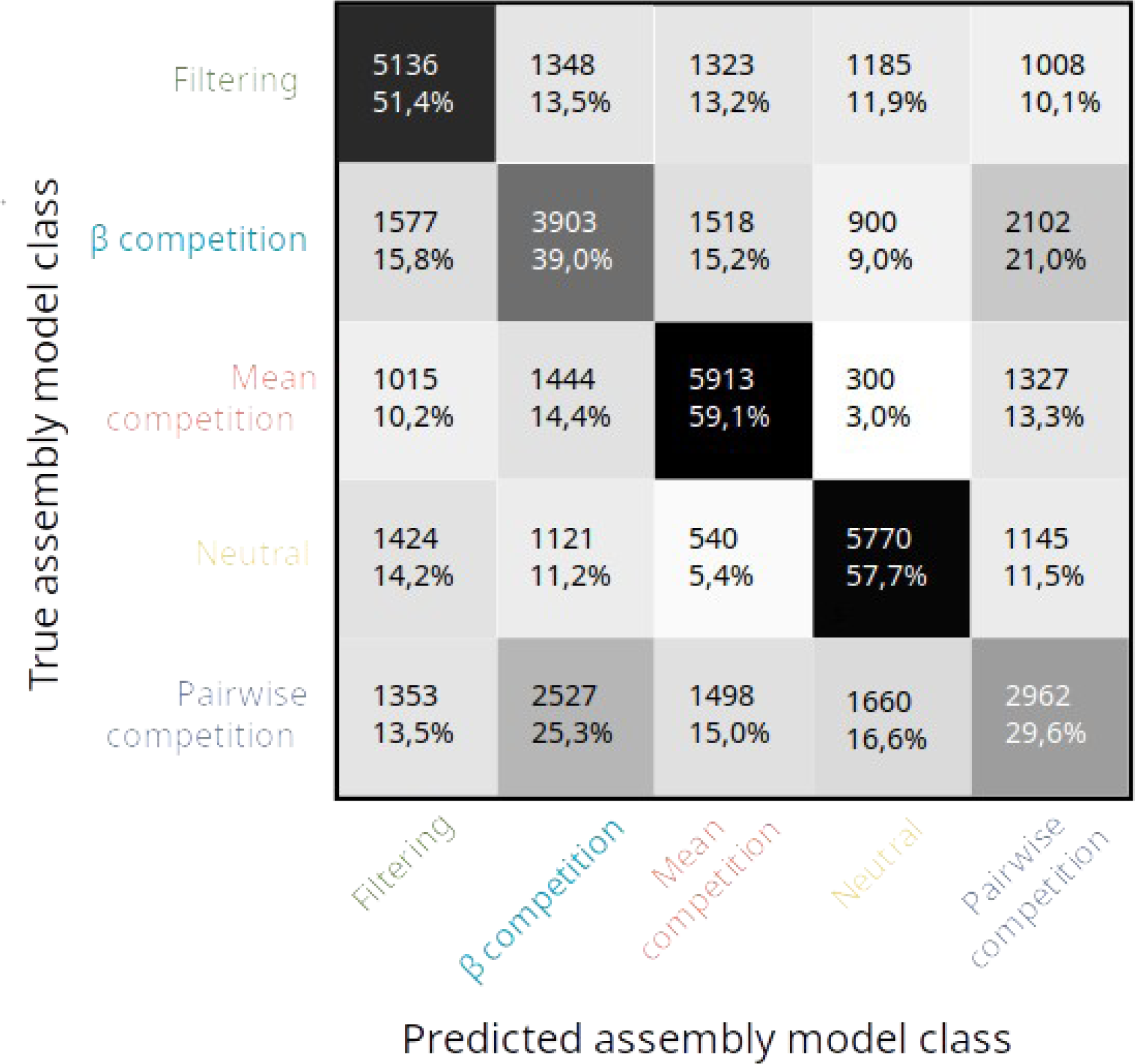
Machine learning classification confusion matrix for datasets simulated under the 5 community assembly models and classified using only trait and genetic diversity data (as is the case for the subtropical forest trees and Galapagos snails datasets). Numbers correspond to the number of datasets simulated under a given community assembly model (rows) that are classified in each model (column). In the case of perfect classification, all values would fall along the diagonal. Percentages indicate the proportion of simulations run with one given class (row) assigned to the column class.

We first consider the three datasets with SAD and genetic data: for the Reunion spider dataset, the confidence percentage in favour of competition is around 40% (Fig. 4) while it was not inferred in [29]. For the two Mascarene weevil datasets, the confidence percentage predicted for the neutral model remains the same as in the analysis by [29], but the circa 40% confidence for both mean competition and filtering in the original analysis is now exceeded by the combination of pairwise competition and *β*-competition (Fig. 4). Pairwise competition and *β*-competition largely dominate over mean competition, which now receives no support. With inclusion of more nuanced competition models, the inference of environmental filtering also now totally disappears in our results for these datasets compared to [29].

**Fig 4.**
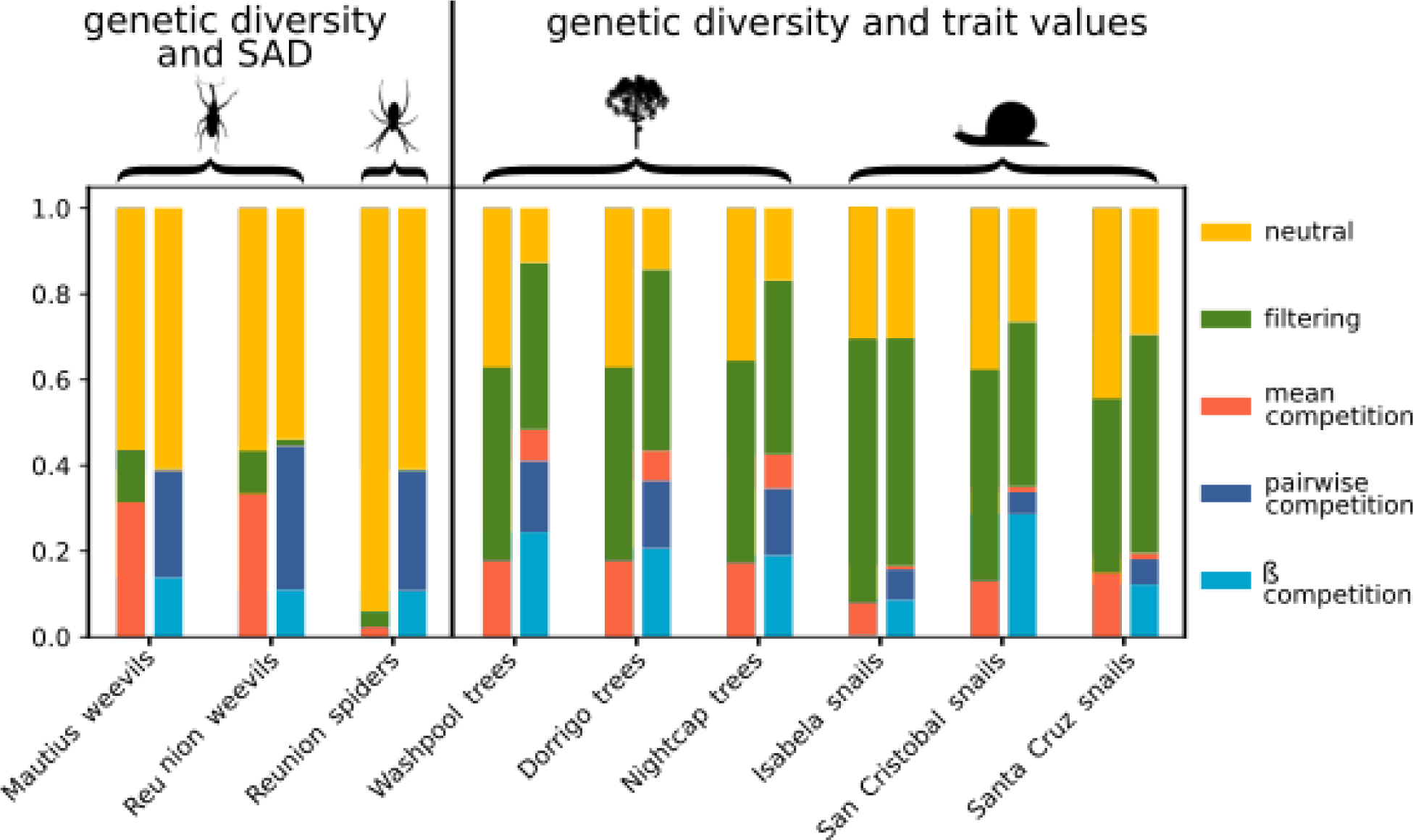
Machine learning classification probabilities for each empirical community for five focal community assembly models. For each dataset, the first bar depicts the result of the original MESS model [29] and the second bar the result with our new competition models. The proportion of colour within each bar represents the proportional predicted model class for neutrality (dark blue), environmental filtering (green), mean competition (orange), pairwise competition (yellow) and *β*-competition (light blue).

Environmental filtering is significantly detected only in the subtropical rainforest tree and Galapagos snail communities, which are also the datasets that contain trait measurements. For empirical data that include trait information, the *β*-competition model was overall a better fit than the other competition models.

For all datasets, the added confidence percentages for all three competition models is exceeds the confidence percentage for competition in [29]. Among competition models, the mean competition is greatly under-represented, and in many cases totally absent when other competition models are available as an alternative. A significant part of the newly identified competition comes from a reduction in the amount predicted for neutrality: data previously predicted to be mainly neutral could be false negatives in the attempt to detect selection (Fig. 4.).

## 3 Discussion

In this manuscript, we investigated the power of inference in community assembly models given different combinations of empirical data. A key advance was the use of a new and more sophisticated competition model that considers the interaction between pairs of individuals instead of making a mean field approximation. Our results show that the mean field approximation can lead to underestimation of the role of competition and overestimation of the role of environmental filtering. We also find that mean competition and environmental filtering produce very similar results in our PCA on approach to equilibrium (Λ < 1) (Fig. 1). This may be because mean competition produces a bimodal trait distribution that is effectively filtering against midpoints in the trait space, while pairwise competition in contrast generates density-dependence mechanisms and allows for a broader range of species to coexist.

In our empirical data analysis, the mean competition model receives almost no support when pairwise and *β*-competition models are added to the analysis as alternatives. This is consistent with the intuition that the new pairwise and *β*-competition models better reflect the biological reality of competition. Indeed, mechanistic simulations with the pairwise competition model were mostly classified by the original MESS inference method [29] as mean competition (S4 Table 3) though sometimes classified as neutral or environmental filtering. This demonstrates that competition can be mistaken for neutrality or environmental filtering if the model of competition is of insufficient complexity. The disappearance of support for the mean competition in our new classifier further supports the hypothesis that pairwise competition is a better description of the empirical data.

Pairwise and *β*-competition simulations have on average more species than the mean competition simulations (see Fig. 2). This is expected because selecting for evenly distributed species across the trait space, as in these competition cases, allows for more diversity than selecting for two diverging groups of species, as in the mean competition case. As *β*-competition depicts density-dependence more accurately, we could expect it to have a significant advantage over pairwise competition. However, the PCA results (Fig. 1) show that simulation outcomes mostly overlap between pairwise and *β*-competition: they could be interpreted as a single indistinguishable category. This is further supported by the confusion matrices (Fig. 6 and S4 Fig. 7 and 8), which suggest that *β_ij_* has no significant influence on the simulation outcome. Density dependence may therefore not play a major role in our analyses of empirical data simply because it was not easily detected by the model selection process. Future work could add further parameters and retrieved summary statistics to better model and better detect density-dependence, but will likely come at a high computational cost.

The striking proximity of the pairwise and *β*-competition simulations to the neutral simulations in our PCA results (Fig. 1) was not apparently consistent with our confusion matrices (Fig. 3 and S4 Fig. 7 and 8). The random forest algorithm seems to be able to distinguish between neutral and non-neutral models, which are indistinguishable for the human eye in the PCA, as well as in most summary statistics (Fig. 2). Weaker inference procedures, backed with less detailed empirical data, may therefore misinterpret competition as neutrality and furthermore, competition-based simulations may often resemble neutral simulations in terms of the community properties studied. This may be an example of emergent neutrality [34], and consistent with niche-neutral models [19] where communities consist of multiple niches but with individuals of multiple species interacting neutrally within each niche. Our results show that despite the now better understood potential for confusion between mechanisms, the combination of ecological data (abundances / traits) and evolutionary (genetic) data, together with machine learning, is a promising approach to distinguish neutrality and selection that outperforms what could be achieved with a single type of data.

The striking difference in our inferences based on the type of data used have implications for the kinds of data we gather to study community assembly. Selection was revealed best by our inference procedure when all data (S4 Fig. 8), or at last trait data, are available (Fig. 3 *versus* S4 Fig. 7). Our result that the neutral model was the best fitting for the spider and weevil datasets that lack trait data seems more likely to be an artefact of data types used in the inference rather than a signal that these communities are assembled by forces closer to neutrality. A comparison of the confusion matrices shows, that while the presence of trait data is not essential for detecting filtering or competition, the more data are available, the better our inference performs (S4 Fig. 8). The signal in the spider and weevil datasets might be too weak to be detected with only 2 data types. Contrary to what has been suggested in the metabarcoding literature [35], our result therefore suggests that genetic data alone may not suffice to measure the selective pressure on a group, traits may be needed as well [30, 36].

We hope that future work will extend our model and results to include macroevolutionary models. If competition is taken into account in existing phenotypic evolution models [37], it often relies only on the species trait value, and does not use abundance data. However, if competitive interactions are mostly pairwise interactions, then the relative abundance of each species would also be relevant when modelling evolution using their trait value, as it would largely influence the strength of intra- and inter-specific competition undergone through density-dependence mechanisms.

During our inference process on empirical data, the selected model is either neutral, competitive (in one of a number of ways) or with environmental filtering. There was not a single model simulated that combines all these processes in varying amounts. Another fruitful direction for future work would be to simulate a simultaneous combination of all the processes in a single model. This would enable us to verify that our inferences (choosing between starkly contrasting models) correspond to what would be predicted by a more nuanced and continuous view of mixed community assembly process. Another direction would be to add intraspecific trait variation which could enable a different handling of the distinction between inter- and intra-specific competition and thus permit several species to occupy the same niche [38]. The *β* factors used in the simulations could also be refined in future work to allow for differences among each pair of species to reflect species-specific interactions, which may generalise to include positive interactions as well as direct competition. This would however necessitate a wide parameter exploration in the simulation, which lead to exponentially greater computational time complexity.

We hope that future empirical studies will be inspired by these findings to provide datasets with all three types of data (genetic diversity, SAD and trait values) rather than having to rely on only two of these three as we did in our present work. As underlined by the supplementary S4 Fig. 8, this would enable our classifier to reach an overall 65% accuracy or 75% accuracy in model classification if we collapse pairwise competition and *β*-competition in a single group, as they are the most similar mechanistically and the hardest to distinguish. Our confusion matrices (Fig. 3 and S4 Fig. 7 and 8) show that the absence of trait data makes it harder to distinguish between the different forms of selection, and future empirical datasets with all three data axes, could be used to verify this sensibility of the prediction to the used data axis.

Our study highlights the importance of the range of empirical data available to detect the ecological footprint of selection, in contrast to neutrality. Our results reiterate a warning that we should not jump too quickly to conclusions about the presence or absence of selection, especially when only one type of data is available. We show that our pairwise competition model (and similar *β*-competition model) are a clear improvement of the previously used mean competition model. Failure to detect pairwise competition in some data sets likely means that competition does not act this way, not that competition, or selection in a broader sense, are absent. We hope that this work will pave the way to improved mechanistic eco-evolutionary models and associated inference procedures for community assembly. We also hope to inspire new empirical data collection and place greater emphasis on the synergistic power of genetic, abundance and trait data when analysed jointly.

## 4 Material & Methods

### 4.1 The MESS model

Our simulations are individual-based with a distinct metacommunity and local or island community [1, 25, 39]. Simulations are run as a time series, enabling the study of both dynamic equilibrium and non-equilibrium behaviour. A single trait value is associated with each species identity, which can be used in different ways to model non-neutral dynamics. After the community simulation is completed and population size fluctuations for each species are known, this information is used to constrain a coalescence-based simulation of genetic variation within each species [40].

Following the MESS model of [29], we simulate a fixed number of individuals in the local community. Each individual *i* is characterized by its trait value *z_i_*. At each time step, one individual dies and is replaced by another individual, which comes either from immigration from the metacommunity, at rate *m,* or from a reproduction event within the local community. We apply selection on the death event because it occurs first though we do not expect to see any meaningful difference from modelling a uniform death process alongside birth rates that are influenced by selection on individual traits. Speciation occurs by point mutation with probability *ν* at each reproduction event. The metacommunity is modelled as a very large regional pool, which is fixed with respect to the timescale of the assembly process in the local community. It arises from ecological and evolutionary processes, including speciation *sensu* Hubbell [1].

Under the assumption of neutrality, the probability of death *P_neutral_* for any given individual *i* in the local community at each time step is given by

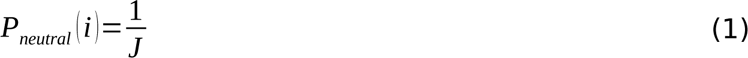

where *J* is the number of individuals in the local community. Selection is incorporated in MESS by computing, at each time step, each individual’s probability of death according to a chosen model of selection (competition or environmental filtering).

In the environmental filtering model, the trait value of each individual is compared to an optimal trait value that depends solely on the environment. The death rate *q_filt_* of any given individual *i* is computed as

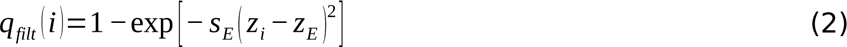

where *z_E_* is the environmental optimum and *s_E_* determines the strength of the filtering. Intraspecific variation is assumed to be negligible in face of interspecific variation, and all individuals of the species *a* have the same trait value *z_a_* which represents the mean phenotype of the species. The probability of death in the next time step, for any given individual is given by the normalized death rate 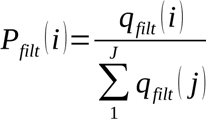

In [29], competition is modelled by a mean-field approximation: the trait value of an individual is compared to the mean trait value of the local community. The death rate *q_MF_* of any given individual *i* is then given by

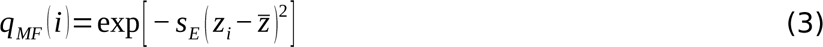

where 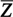 is the local community mean trait and s*_E_* determines how quickly competitive pressure decays with the distance between trait values. Just as for environmental filtering, the death probability *P_MF_* (*i*) for each individual *i* is derived through normalization by

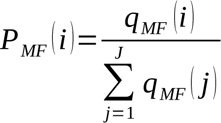

The mean-field approach collapses all trait differences into one value and can therefore generate counter-intuitive results. For example, the distribution of species across the trait axis might be bimodal as two groups of species diverge away from the central mean value, leading to an obvious gap around the mean (see S3 Fig. 6). The area around the mean in trait space is thus free from species and competition but is still the most penalised trait, while denser areas, further away from the mean but with more species, are favoured.

Here, we correct this artefact by using a new and more realistic competition model based on pairwise comparisons between all individuals. In our model, the death rate *q_pair_* of any given individual *i* is based on the mean of all pairwise trait differences with the other individuals in the local community:

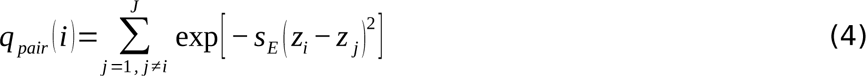

The added computational cost of the pairwise model was partially offset by optimizing the underlying data structures of the original MESS model [29], which was essential due to the large number of simulations needed to train our inference procedure (see S2 Table 2). In contrast to the mean competition model, the pairwise competition model is expected to produce uniformly and regularly distributed species along the trait axis, which is confirmed by test simulations (see S3 Fig. 6). The pairwise competition model does not, however, allow us to refine the strength of intra-specific competition: individuals of the same species have the exact same trait value and thus the exponential in equation (4) is always equal to 1. To allow investigation of this, we also implement a third “*β*-competition” model that introduces an interaction matrix parameter *β_ij_* to modulate competition strength between all possible pairs of individuals. Larger values of *β_ij_* increase the strength of competition between individuals *i* and *j*. We set *β_ij_* = *β*_intra_ when individuals with indexes *i* and *j* are conspecific, and *β_ij_* = *β*_inter_ when they are heterospecific. The resulting death rate is given by

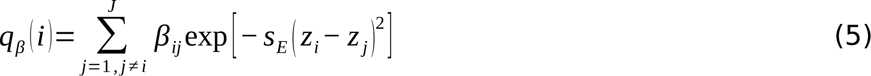

By allowing intra- and inter-specific competition to differ according to a parameter, we are in effect modelling differing levels of negative density dependence: *β*_intra_ >> *β*_inter_ corresponds to strong intraspecific density dependence whilst *β*_intra_ << *β*_inter_ corresponds to no density dependence. We leave the *β*_intra_ << *β*_inter_ case for future work, noting that preliminary tests suggest the model will lead to mono dominance. The three competition models that we study here are summarised in Fig. 5. Notably, the death probability for each individual, computed from the given death rates, converges toward a neutral probability 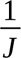 as the strength of selection *s_E_* converges toward 0, in accord with the theory of a continuum spectrum from neutrality to strong selection [6].

**Fig 5.**
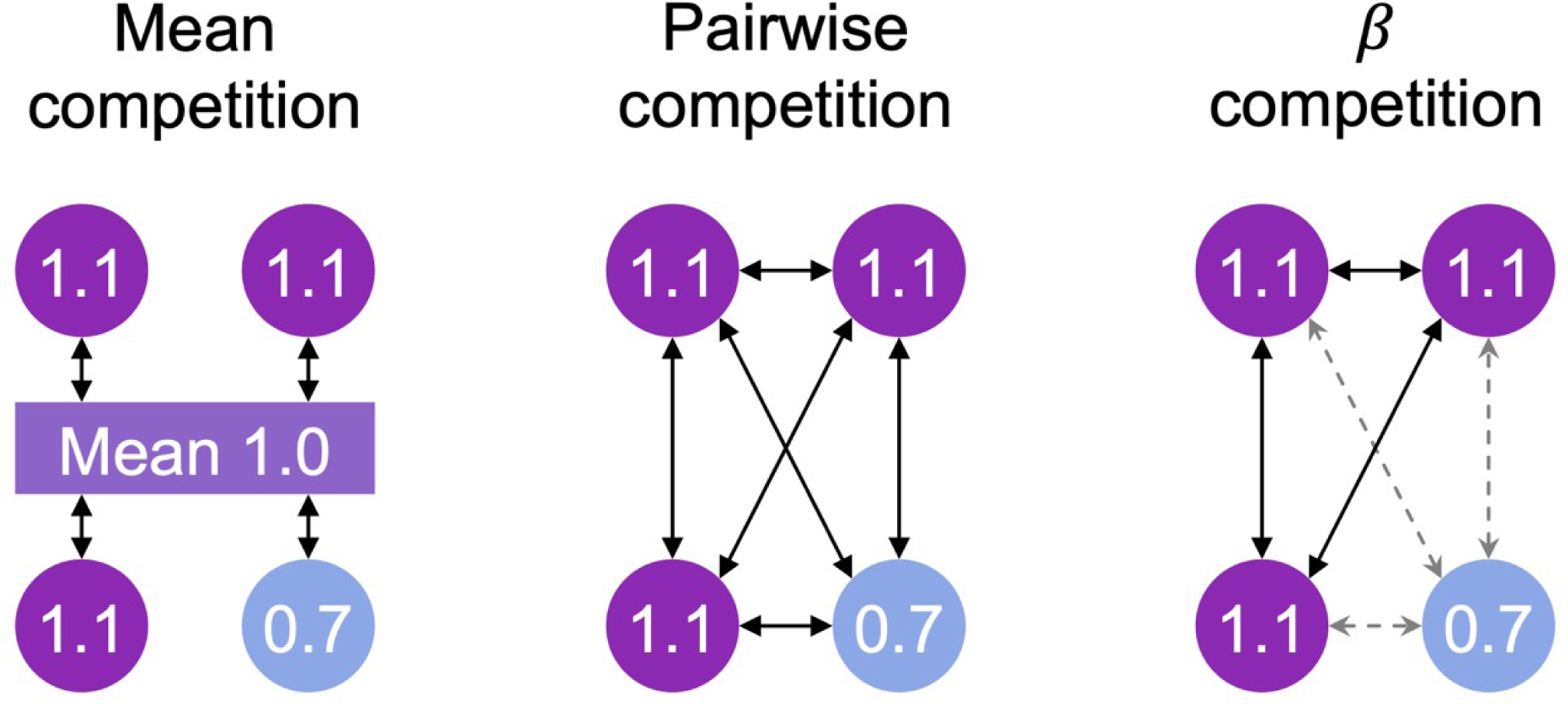
Depiction of the different forms of competition. Each circle represents an individual, with the colour specifying which species it belongs to and the number its (one dimensional) trait value. The effect of competition on fitness (symbolized by arrows) is shown for all individuals. Mean competition model: the trait value of each individual is compared to the mean trait value for the community to which all species contribute. Pairwise competition: the trait value of every individual is compared individually to every other individual’s trait value. *β*-competition: the trait value of each individual is compared individually to each other individual’s trait value, weighted by a factor depending on whether the pair of individuals belong to the same species. The style of arrows in the case of *β*-competition symbolizes the type of competition: inter-specific competition (solid black arrows) or intra-specific competition (dotted gray arrows).

### 4.2 Exploration of *in silico* experiments

To explore the behaviour of the proposed competition models and understand how the different models affect the outcome of community assembly, we ran 10 000 simulations for each of the five community assembly models (neutral, filtering, mean competition, pairwise competition and *β*-competition), covering wide ranges of possibilities for the main parameters of the simulations : the age of the community (through Λ, a parameter used to quantify the progress of the simulation toward equilibrium), the number of individuals *J*, the strength of the ecological filtering or competition *s_E_*, the strength of inter-individuals interactions *β*, the migration rate *m*, the speciation rate *ν*, and the abundance/effective population scaling factor *α* (see S1 Table 1).

We compare our results to those from the previous implementation of MESS to illustrate the important effects of our improvement. To do this, we use the same simulation descriptors as [29]. To briefly summarise these here, each simulation is characterised by a number of summary statistics along each data axis (species abundances, population genetic variation and trait values). These summary statistics are: the first moments of each community-wide distribution, Spearman rank correlations among all data axes, differences between metacommunity and local community values of trait mean and standard deviation, and Hill numbers of several orders to quantify the shape of each distribution [41]. Hill number of order *q* for a data axis *X* (SAD, traits data or genetic diversity data), will be noted ^q^X. These calculations were done with built-in functions of MESS, and the detailed method is described in the supplementary material of [29]. The temporal trends are studied in terms of Λ, a parameter used to quantify the progress of the simulation toward equilibrium [25], used in common with the original MESS model [29]. A community is considered at equilibrium, and Λ=1, when the initial conditions are no longer detectable in the system, and this advancement toward equilibrium is measured as the proportion of individuals in the community descending from a lineage that colonized during the simulation [39]. We visually inspected the resulting simulations by collapsing simulated summary statistics using a PCA after [29] (Fig. 1). This enabled us to distinguish between the different community assembly models.

### 4.3 Machine learning and inference

We follow the same procedure as [29] for model classification and parameter estimation: Random Forest [42] with python and the scikit-learn module (v0.20.3) [43]. We first train a machine learning classifier in a supervised fashion on 50,000 simulated datasets (10,000 for each assembly model). We then use the trained classifier to predict model class probabilities for each of the empirical datasets. A confidence percentage is associated to each model. We quantified classifier accuracy using 5-fold cross-validation on simulated data and evaluated model misclassification by combining these results into a confusion matrix. We evaluated classifier accuracy using three different suites of simulated data axes, one composed of SAD and genetic data, another composed of trait values and genetic data and a third corresponding to an ideal case scenario, with all three data axis. The first two of these simulated data sets mirror the data configurations of our empirical datasets. Results from the third data configuration demonstrate that extensive gathering of empirical data would substantially improve the performance of the classifier (see S4 Fig. 8).

### 4.4 Study of empirical datasets

We used the empirical datasets following [29]: 1) a spider community from Réunion island with standardized sampling for abundance and genetic diversity of ten 50 m x 50 m plots and 1282 individuals sequenced for one 500bp mtDNA region (COI) [44]; 2) two weevil communities from two Mascarene islands (one from Réunion and one from Mauritius) which have been densely sampled for abundance and sequenced for one mtDNA region (600bp COI) at the community-scale [45]; 3) three subtropical rain forest tree communities scored for multiple continuous traits and shotgun sequenced for whole cpDNA [46]; 4) Galapagos snail communities collected from all major islands (three in total), sampled for one mtDNA region (500bp COI; [47]) and scored for two continuous traits [48]. We compared summary statistics linked to the SAD, genetic diversity and traits computed on the empirical data to those computed on 50,000 simulations (10,000 for each community assembly model).

## Acknowledgments

This study was enabled through the financial help of the École Normale Supérieure -- PSL. Much of the original research was conducted by JL during a placement to Imperial College with JR. We thank both the Imperial College research computing services (High Performance Computing) and the IBENS for the usage of their computation cluster. Through JR, this study is an output of the Georgina Mace Centre for the Living Planet at Imperial College London.

## Notes

### Competing Interest Statement

The authors have declared no competing interest.

### Summary of Updates

Main text clarified, figures reorganized and updated

