## Supplementary Material for "Detecting the ecological footprint of selection"

#### S1 Model's parameters

| Categorical parameters | | | |
|---|---|---|---|
| Parameter | | Options | Tested range |
| Community assembly model | | Neutral / Competition / Environmental filtering / Pairwise competition / $\beta$-competition | All options † |
| *In situ* speciation model | | None / Point mutation / Random fission | Point mutation |
| Local community initial conditions | | Metacommunity sample / Monodominance | Monodominance |
| **Metacommunity component parameters** | | | |
| **Symbol** | **Meaning of parameter** | **Type and range** | **Tested range** |
| $J_M$ | Total number of individuals | Integer 1 | 5e5 |
| $S_M$ | Total number of species | Integer > 1 | 250 |
| $\lambda$ | Per lineage birth rate (speciation) | Real ∈ [0,∞] | 2 |
| ε | Per lineage death rate (extinction) as proportion of $\lambda$ | Real ∈ [0, 1] | 0.7 |
| $\sigma^2 M$ | Trait evolution rate variance (Brownian motion) | Real > 0 | 2 |
| **Local community component parameters** | | | |
| **Symbol** | **Meaning of parameter** | **Type and range** | **Tested range** |
| $J$ | Total number of individuals | Integer > 1 | [1000, 5000] † |
| $S$ | Local species richness* | Integer > 1 | *Not applicable* |
| $\nu$ | Per capita per birth speciation rate | Real ∈ [0, 1] | [0.0005, 0.005] ° |
| $m$ | Immigration rate from metacommunity (per step) | Real ∈ [0, 1] | [0.001, 0.01] ° |
| $\sigma^2$ | Trait evolution rate variance (per speciation event)* | Real > 0 | *Not applicable* |
| $z_E$ | Optimal trait value in environment* | Real | *Not applicable* |
| $s_E$ | Strength of ecological filtering | Real > 0 | [0.01, 10] ° |
| $\Lambda$ | Fraction of turnover equilibrium* | Real ∈ [0, 1] | [0, 1] † |
| $\beta_{intra}$ | Strength of intraspecific competition | Real > 0 | [0.01, 1] ° |
| $\beta_{inter}$ | Strength of interspecific competition | Real > 0 | [0.01, 1] ° |
| **Population genetics coalescence component parameters** | | | |
| **Symbol** | **Meaning of parameter** | **Type and range** | **Tested range** |
| $L$ | Sequence length of simulated genomic region | Integer > 0 | 570 |
| $\mu$ | Mutation rate | Real ∈ [0, 1] | 2.2e-8 |
| $\alpha$ | Abundance/Ne scaling factor | Integer > 0 | [1000, 10000] † |

**Table 1: MESS model parameters**

All MESS model parameters, their interpretations and range of possible values.

Parameters indicated with an asterisk (*) are pseudo-parameters which are either emergent, compound, or randomly sampled from a distribution with parameters determined by other elements of the model.

Parameters for the simulations where either uniformly (†) or loguniformly (°) drawn in the range referenced as tested range, when applicable. The chosen ranges are based on [29].

#### S2 Algorithm optimisation

A pairwise comparison of trait values costs more computation time than a single comparison to the mean. As expected, the simulations for the pairwise competition model were slower (around 4 times) than for the mean competition model. Yet, the computation time is a critical issue in a context of large numbers of simulation experiments required for classifier training, and this necessitated code optimization.

| **Model** | **Before optimization: [29] with pairwise competition added by this study** | **After optimization: this study** | **Ratio** |
|---|---|---|---|
| **Neutral** | 9 ± 2 | 24 ± 3 | 0.375 |
| **Mean competition** | 456 ± 114 | 89 ± 11 | 5.14 |
| **Pairwise competition** | 1944 ± 258 | 323 ± 40 | 6.02 |
| **Environmental filtering** | 80 ± 18 | 70 ± 10 | 1.14 |

**Table 2: Comparison of the speed of the simulations for different version of the code** (mean value per run, ± standard deviation, in seconds). Results are for 50 simulations of 200 generations on a single core.

### S3 Trait distribution exploration

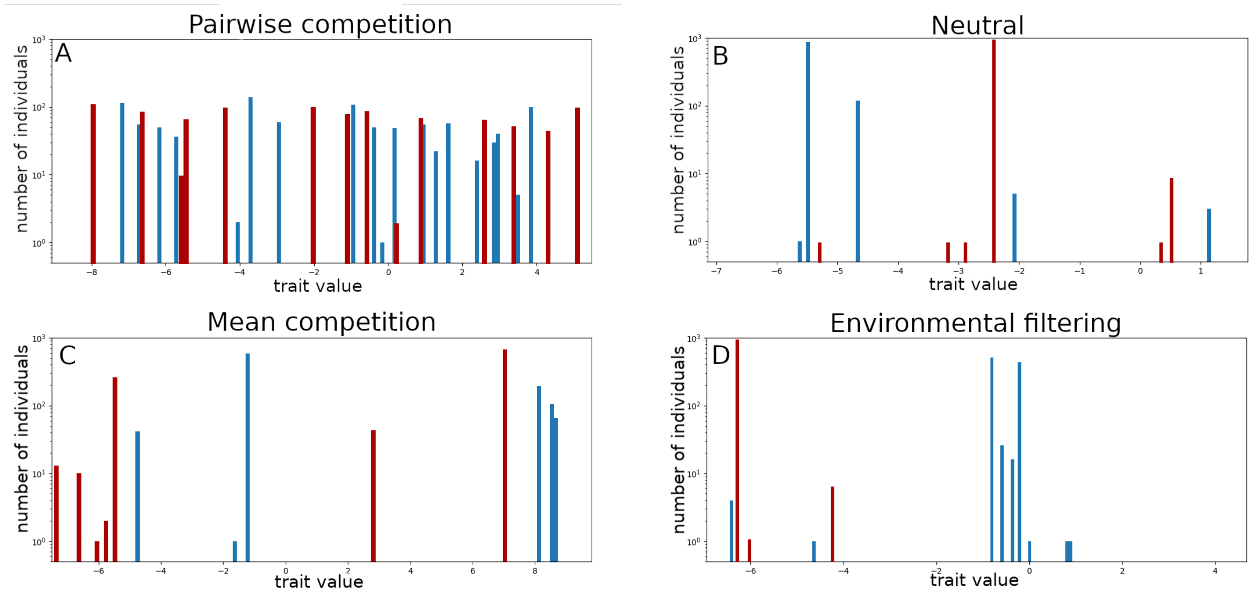


**Figure 6: Typical trait values distribution for four studied 4 community assembly models. Two examples (red and blue) are given for each model.** Two groups of species are distancing themselves in the mean competition model (C), while the species are much more grouped together in the environmental filtering case (D) and evenly distributed in the pairwise competition model (A). In the neutral case, they are random and their abundances follow a typical log-normal distribution. This also shows that we can expect significantly different results in the summary statistics resulting from trait data, but also in the species abundances and their variation and thus in the phylogeny.

#### S4 Machine learning assessment

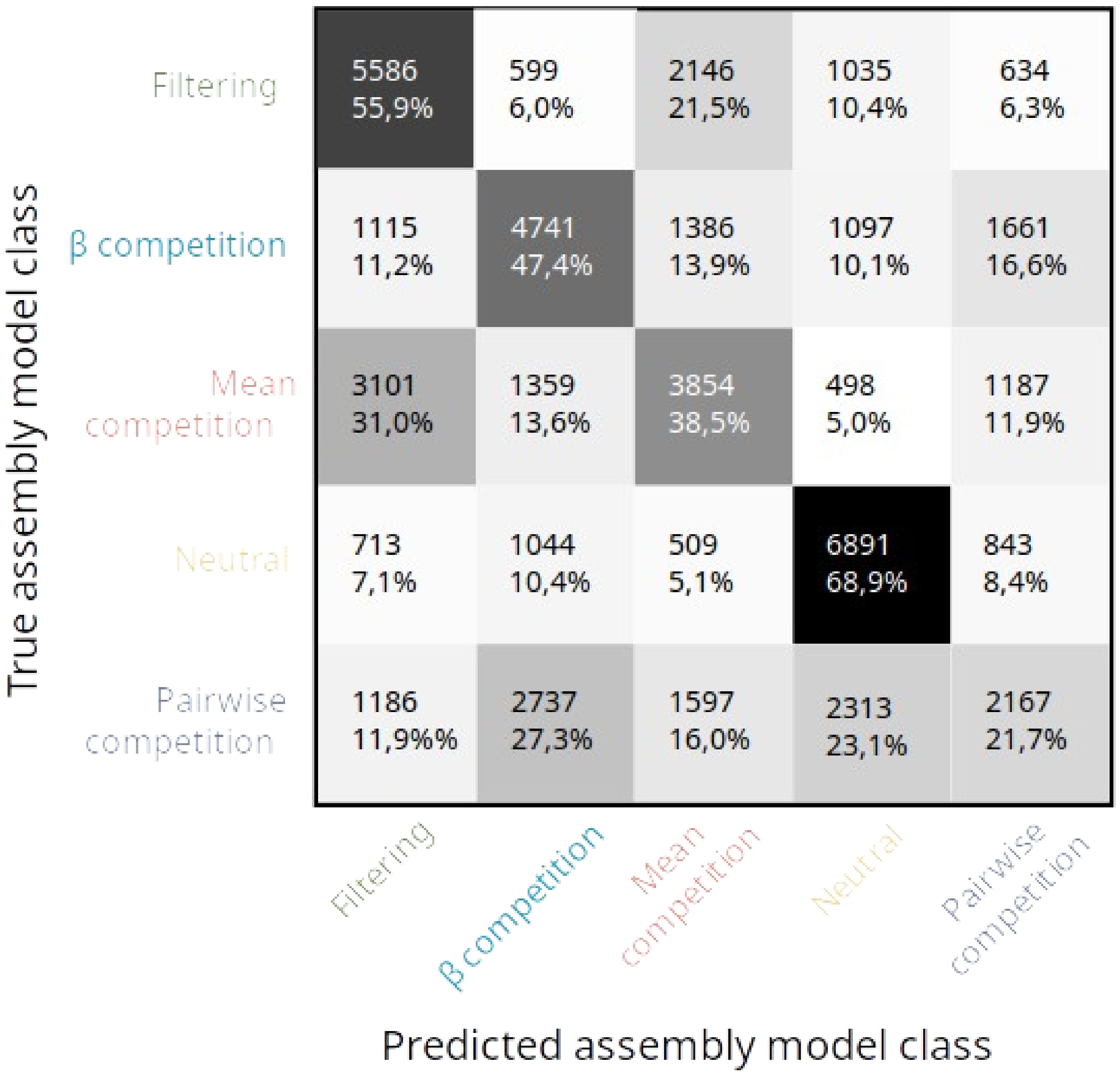


**Figure 7: Machine learning confusion matrix for data set produced by simulation using the 5 community assembly models and classified using only SAD and genetic diversity data**. Percentages indicate the proportion of simulations run with one given class (raw) assigned to the column class. Mean competition is often mistaken for filtering, and pairwise competition for both neutrality and $\beta$-competition.

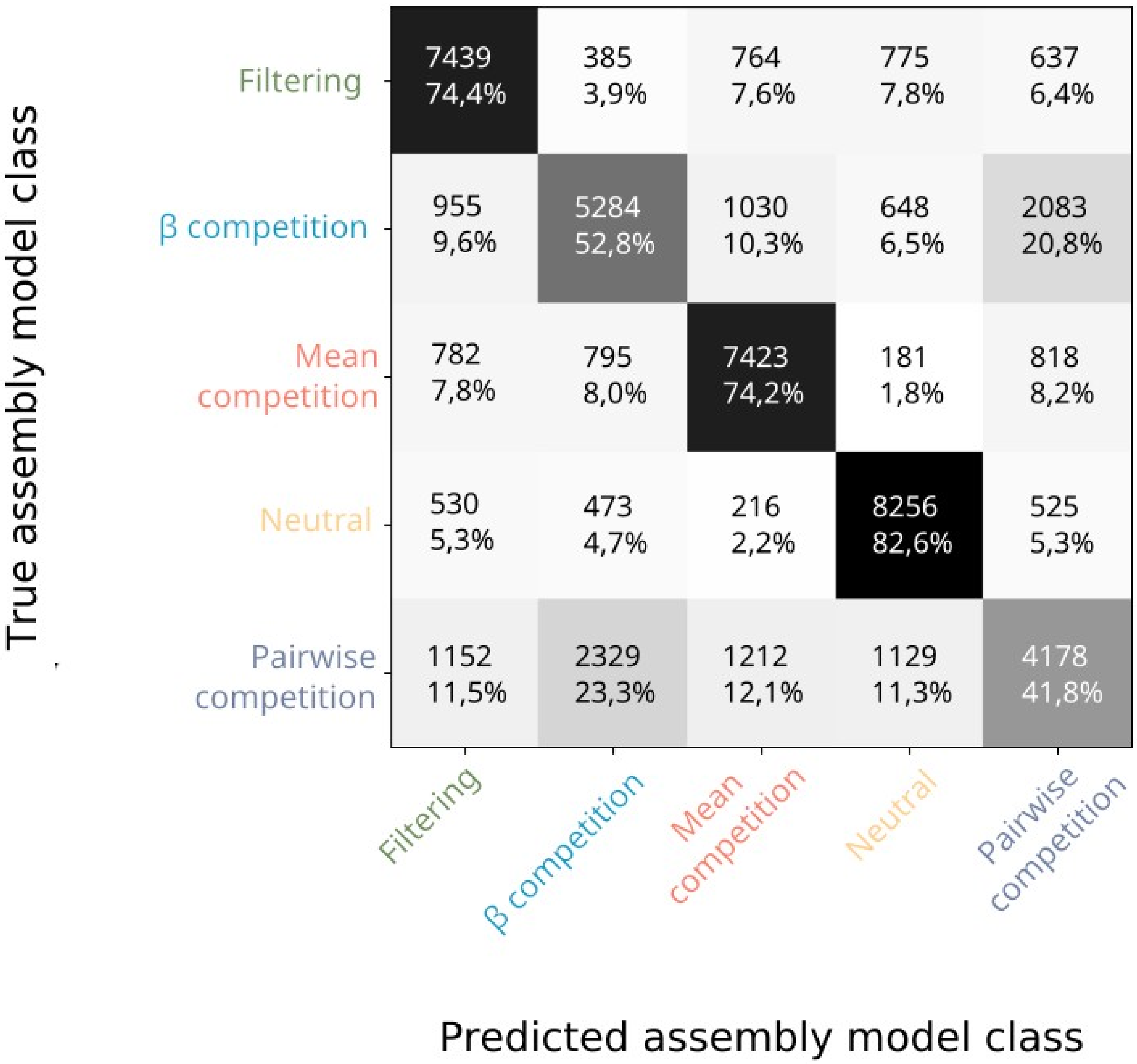


**Figure 8: Machine learning confusion matrix for data set produced by simulation using the 5 community assembly models and classified using all three data axis (SAD, trait and genetic diversity).** Percentages indicate the proportion of simulations run with one given class (row) assigned to the column class. 80% of the time, neutral simulations are correctly distinguished from non-neutral simulations.

| Model | Mean competition | Neutral | Filtering |
|---|---|---|---|
| Percentage of pairwise simulations assigned to the model | 48% | 28% | 24% |

**Table 3:** Inference of 100 simulations run under the pairwise competition model classified by a classifier trained only with mean competition, neutral and environmental filtering simulations.
